# cFAR and Relative Signal: Diagnosing Preferred Orientation in Single-Particle Cryo-EM

**DOI:** 10.64898/2026.08.11.744264

**Authors:** Valentin Peretroukhin, Michael McLean, Ali Punjani

## Abstract

The quality of single particle cryo-EM reconstructions can be severely degraded when an insufficient variety of 3D particle orientations is present in the image data, limiting downstream model building and interpretation. However, it is often difficult to ascertain whether or not a particular dataset suffers from such preferred orientation since the required orientation coverage depends on target geometry, alignment accuracy, and particle quality. To simplify diagnosis of preferred orientation, we present two complementary methods. First, the conical Fourier Shell Correlation Area Ratio (**cFAR**) compares the worst- and best-correlating conical regions of 3D Fourier space to quantify half-map anisotropy into a single, easily interpretable score ranging from zero to one. Second, **Relative Signal**, a companion to cFAR, directly relates signal content to viewing direction so that under-sampled views can be identified. We characterize our methods and compare them to existing anisotropy detection approaches on synthetic data and on 14 real datasets that span sundry molecular weights and structure types. Implementations of both cFAR and Relative Signal are included in CryoSPARC v4.5 and later versions.

## 1. INTRODUCTION

Single-particle electron cryomicroscopy (cryo-EM) aims to reconstruct a high-resolution three-dimensional structure of a biological macromolecule (the “target”) from two-dimensional images of frozen copies (“particles”) of that molecule. To accomplish this, a purified sample containing the target molecules is prepared and applied to a mesh grid, which is rapidly flash-frozen with the goal of producing a thin layer of amorphous ice in which the target molecules are ideally randomly oriented. The grid is then placed in a transmission electron microscope and different parts of the grid are imaged to capture as many copies of the target in as many different viewing directions as possible. After data collection, commonly-used analysis software (e.g., CryoSPARC^1^, RELION^2^, cisTEM^3^) is used to identify, extract, and curate a set of particle images that can then be used to generate a three-dimensional reconstruction. It is vital to collect and curate a diverse set of viewing directions of the target for reconstruction algorithms to yield high-quality structures.

In many realistic scenarios, however, a set of particle images will not contain a sufficient sampling of orientations of the target. Particles may, for instance, adopt “preferred orientations” due to air-water interface effects or a tendency to adhere to the supporting grid^4^, and therefore any set of collected images will inherit this bias. The set may also mix conformational or compositional states of the target, along with “junk” outlier images that do not contain the target at all, complicating the reconstruction process. Finally, particle picking and curation methods themselves may introduce anisotropic bias even if the ice contains a sufficient variety of orientations (e.g., certain views of the target may be more difficult to identify or align). In any of these cases, reconstructing a 3D density map from a deficient set of views can yield severely degraded map quality characterized by distinctive smearing, stretching, or squeezing artifacts (e.g., densities in Fig. 1). In practice, identifying preferred orientation is non-trivial and a careful visual inspection of the final reconstruction by a human expert is often necessary.

**Figure 1.**
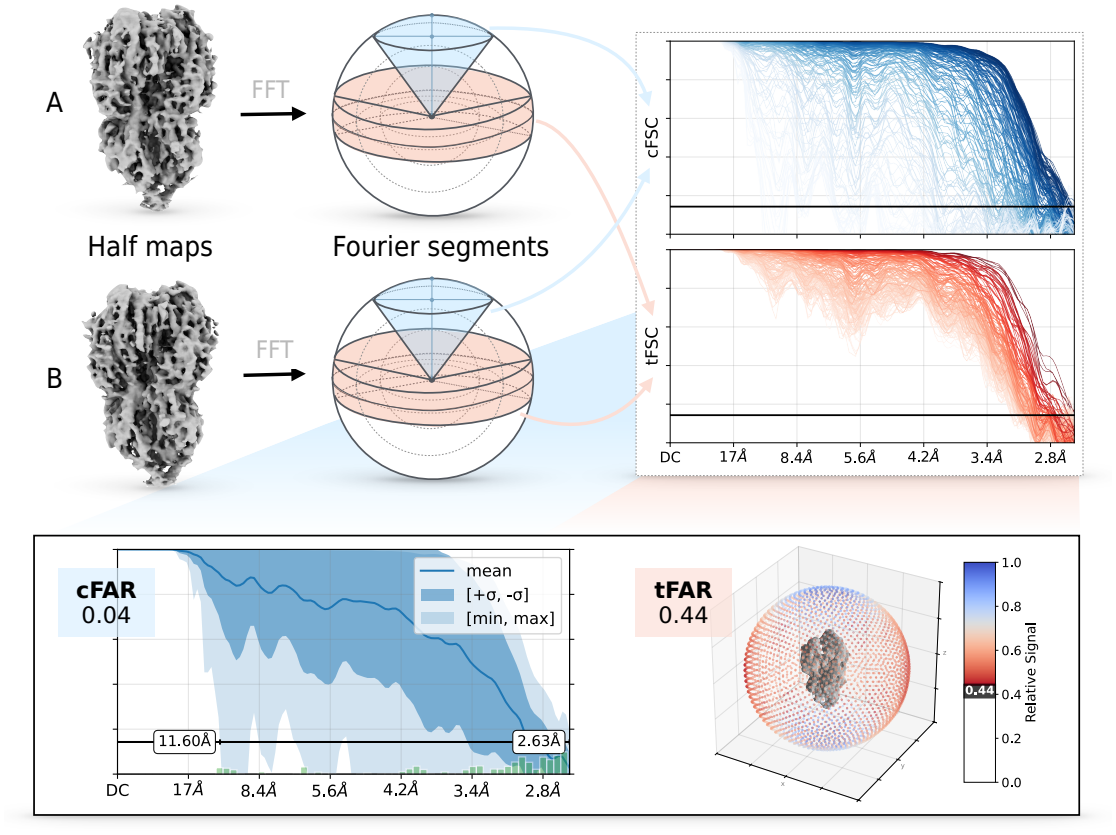
Our approach to diagnosing preferred orientation. We compute Fourier shell correlations of two half-maps within a set of conical and toroidal segments of the Fourier sphere (cFSCs and tFSCs, respectively). Given these two sets, we compute two ratios, `cFAR` and `tFAR`, that aim to identify preferred orientation by comparing extrema. We summarize cFSC curves with per-shell statistics and 0.143 crossings (bottom left). We define a normalized area under tFSCs as Relative Signal and visualize it on the viewing sphere of the consensus density (bottom right).

To aid and expedite this process, we present two numerical approaches (included in CryoSPARC ≥v4.5) that can quickly and effectively diagnose the presence of preferred orientation. Namely:

1. The conical Fourier Shell Correlation Area Ratio (`cFAR`): a metric comparing the worst-to the best-correlating conical segment of 3D Fourier space and yielding a single, easily interpretable number ranging from zero (severe anisotropy) to one (uniform resolution among all directions).
2. The toroidal Fourier Shell Correlation Area Ratio (`tFAR`), an analogous ratio that compares “toroidal” segments, and the related *Relative Signal*, a weighted area under toroidal correlation curves that serves to map directional signal content to viewing direction.

`cFAR` captures the effects of both particle orientation distribution and the distribution of coherent signal in a 3D map, and is therefore able to correctly identify anisotropy in the presence of junk particles which have been assigned incorrect orientations. It is a conservative diagnostic, and is sensitive to the presence of even a single poorly correlated conical region, ensuring that 3D maps that need further inspection are consistently assigned a low `cFAR` score. `tFAR` serves as a less conservative metric, owing to toroidal segments having greater dispersion in Fourier space. Finally, Relative Signal directly identifies *which viewing directions* are underrepresented or are of poor-quality in a dataset, information that can be used to improve upstream particle curation or data collection strategies. `cFAR` and Relative Signal are available in CryoSPARC ≥v4.5 and are very fast to compute, up to an order of magnitude faster than other methods.

To demonstrate these advantages, we compare our methods to four other existing anisotropy metrics in the literature across synthetic data tests and 14 reconstructions from EMPIAR^5^ depositions of real cryo-EM data.

## 2. BACKGROUND

The problem of preferred orientation and its effects on 3D reconstruction have been known in the literature since the development of *negative stain imaging*^6^, a precursor to cryo-EM. In some cases, it is possible to modify sample preparation conditions (typically using detergents) to avoid preferred orientation in a sample^7^. For a sample that has already been vitrified, two primary avenues exist to address preferred orientation:

1. tilt the microscope stage during sample collection^8^ to collect more oblique views of the target if the bias was introduced during sample preparation, and/or
2. rely on specialized picking, curation, denoising, or interpolation methods^9–13^ to find a more varied set of particle images or to correct for data processing bias.

A crucial prerequisite to applying any mitigation strategy, however, is accurately diagnosing the presence and severity of preferred orientation. This is made challenging for a number of reasons. For example, when inspecting a three-dimensional reconstruction, one might consider consulting a histogram of the viewing angles which were recovered by the algorithm to ascertain whether there is sufficient coverage of the viewing sphere. However:

- reconstruction algorithms may assign poor-quality or “junk” images to specific regions of the viewing sphere, and
- it is not always clear what constitutes adequate “coverage”, since this will depend on the quality of the particle images and the target (pseudo-)symmetry.

Indeed, as mentioned in the introduction, a careful visual inspection of the final reconstruction by a human expert is often a necessary and crucial step of diagnosis. Several existing approaches have been explored to attempt to automatically identify the presence of preferred orientation; in this section, we recap the canonical cryo-EM reconstruction procedure and then introduce the existing methods that motivate our approach.

### 2.1. Cryo-EM Reconstruction

The cryo-EM imaging process can be modeled via the forward model:

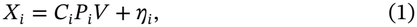

where *V* : ℝ^3^ → ℝ denotes the 3D electrostatic potential of the molecule under study, and *X*_*i*_ ∈ ℝ^*N*×*N*^ is the *i*-th observed particle image. operator *P*_*i*_ denotes the tomographic projection of *V* under pose *T*_*i*_ ∈ *SO*(3) × ℝ^2^ (comprising an unknown 3D rotation *R*_*i*_ and 2D in-plane translation *t*_*i*_), yielding a 2D projection image of the volume as seen from that viewing direction. The operator *C*_*i*_ denotes the contrast transfer function (CTF) and *η*_*i*_ denotes zero-mean additive Gaussian noise.

Typically, an “ab-initio” process will produce an initial estimate of *V* and a CTF estimation procedure will estimate *C*_*i*_, both of which are then used to initialize an expectation-maximization (EM) procedure that iteratively refines an estimate of *V* while simultaneously solving for the unknown poses *T*_*i*_ of each particle image. To guard against overfitting and to provide an unbiased estimate of resolution, this refinement is typically carried out in two independent, parallel tracks, each using a randomly assigned half of the particle stack; this produces two independent volume estimates, or “half-maps,” *Va* and *V*_*b*_.

An FSC (Fourier shell correlation)^14^ can be computed between *Va* and *V*_*b*_ to assess the resolution and self-consistency of the reconstruction. For a given spatial frequency *r*, the FSC is defined as the normalized cross-correlation between the two half-map Fourier transforms, restricted to the spherical shell of radius *r*:

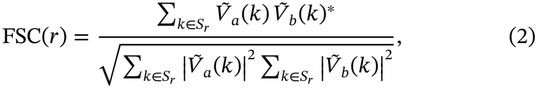

where 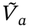 and 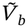 denote the 3D Fourier transforms of the two half-maps, *S*_*r*_ = {*k* ∈ ℝ^3^ : ||*k*|| = *r*} is the set of Fourier voxels lying on the shell of radius *r*, and (·)* denotes the complex conjugate. The Gold-Standard FSC (GSFSC)^15^,^16^ is defined as the FSC curve computed between two half-maps that have been refined entirely independently from the outset — including independent particle assignment, independent pose/CTF estimation, and independent reconstruction — so that the two maps share no information beyond the initial model. The nominal resolution of the reconstruction is then reported as the spatial frequency *r*\* at which FSC(*r*) first crosses below a fixed threshold, conventionally 0.143, converted to the reciprocal units of Å.

### 2.2. Existing Preferred Orientation Metrics

Given *Va, V*_*b*_, and the set of particle orientations {*R*_*t*_}, there are several existing approaches to detecting potential preferred orientation. We describe four approaches here that we will later use to compare against our work.

#### 2.2.1. *Efficiency of Orientation Distribution (E*_od_)

We start with Naydenova and Russo^17^ who introduced a method called cryoEF. This framework relies on the fact that every orientation distribution has an associated point spread function (PSF) in real space. The effective resolution in a given direction depends on how many views contribute information along that direction, and an anisotropic distribution of viewing angles produces a PSF that is elongated along poorly sampled directions and narrow along well-sampled ones. The PSF radius 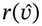, sampled over all directions 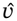, thus serves as a proxy for directional resolution: large 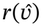 indicates poor resolution along 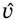, while small 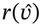 indicates good resolution. Letting 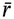 and *σ* denote the mean and standard deviation of 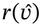 across directions, Naydenova and Russo define the “efficiency” of an orientation distribution, *E*_od_, as:

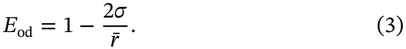

**Notes on** *E*_*od*_

A perfectly uniform distribution has *σ* = 0 ⇒ *E*_od_ = 1 (isotropic PSF); in extreme cases *E*_od_ can fall slightly below 0.

#### 2.2.2. *Sphericity of 3D Fourier Shell Correlation (*Ψ)

Next, Tan et al.^8^ introduced the directional FSC, which generalizes the ordinary (global) FSC by restricting the comparison between half-maps to a narrow cone of Fourier directions rather than pooling over the full shell. For a cone of directions about axis 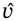, the FSC is computed between half-maps restricted to that cone, denoted 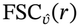. Repeating this over many axes and interpolating onto a grid yields a full 3D volume of directional resolution values, the 3DFSC. The per-direction resolution is defined as the 0.143 crossing of 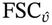, exactly as for the ordinary global FSC, but now evaluated independently for each direction 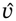.

Since the 3DFSC assigns a resolution value to every direction, it can be visualized as a 3D surface: directions with better resolution sit farther from the origin, and directions with worse resolution sit closer in. For an isotropic reconstruction, this surface should be (close to) a perfect sphere; anisotropy distorts it into a flattened or lobed shape. To quantify this, the 3DFSC is thresholded (here, we choose to threshold at 0.5) to obtain a closed iso-surface enclosing a volume *V*_*p*_ with surface area *A*_*p*_. The “sphericity” of this surface provides a metric for preferred orientation and is defined by the Wadell measure, originally developed in sedimentology to quantify how closely grain shape approximates a sphere^18^:

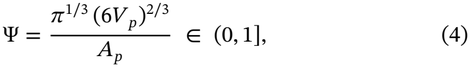

**Notes on** Ψ

Ψ ≤ 1, Ψ = 1 for a perfect sphere (isotropic resolution), with Ψ ≳ 0.8 commonly taken to indicate near-isotropic resolution.

#### 2.2.3. Sample Compensation Factor (SCF)

Baldwin and Lyumkis^19,20^ introduced the Sample Compensation Factor (SCF) as a measure of preferred orientation. SCF is computed from the geometry of ‘Fourier sampling,’ rather than from any measured correlation between half-maps. By the Fourier slice theorem, a particle with viewing direction 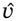 populates a thin slab of Fourier space about the central plane orthogonal to 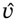. On the shell of radius *r*, the voxel at 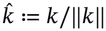 is thus sampled by that particle if and only if

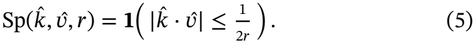

Summing over all particle views yields the accumulated per-bin sampling 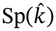 on each shell.

With ⟨⋅⟩ denoting the arithmetic mean over all bins of a shell,

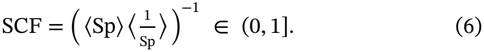

This quantity is exactly 1 for uniform sampling and shrinks toward 0 as sampling grows more uneven; it is undefined if any bin has zero sampling. To remain well-defined when some bins are entirely unsampled, let *q* denote the fraction of zero-sampled bins, *p* = 1 − *q*, and Sp^*^ the sampling values restricted to nonzero bins:

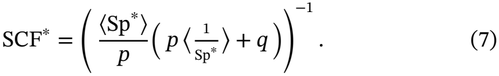

**Notes on SCF**

When *q* = 0, SCF^*^ = SCF. A value of 8/π^2^ ≈ 0.81 corresponds to a ‘perfect side distribution,’ and values above this threshold generally indicate adequate sampling^20^.

#### 2.2.4. Fourier Shell Occupancy (FSO) and the Anisotropy Transition Zone

Finally, Vilas and Tagare^21^ introduced the Fourier Shell Occupancy (FSO) summary plot, built from a set of directional FSCs computed over *N* cone directions. At each frequency *f*, FSO asks a comparative question: what fraction of directions are performing at least as well as the map’s global (isotropic) FSC curve? Letting FSC_global_ denote the ordinary global FSC,

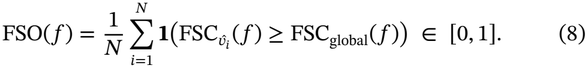

At low frequencies FSO(*f*) ≈ 1, since essentially all directions are well resolved; as *f* increases, poorly-sampled directions fall behind the global average one by one, so FSO(*f*) decreases toward 0. A sharp decline concentrated at one frequency indicates a fairly isotropic map, while a gradual decline spread over a wide frequency range indicates strong anisotropy.

This motivates the *anisotropy transition zone*: the resolution band over which the FSO falls from 0.9 to 0.1. Let *f*_0.9_ and *f*_0.1_ denote the frequencies at which FSO crosses these values (via linear interpolation), with corresponding resolutions *d*_0.9_ ≥ *d*_0.1_; *d*_0.9_ is the resolution up to which the map is essentially isotropic, and *d*_0.1_ is the resolution beyond which nearly all directions have fallen below the global FSC curve. To generate a single numerical quantity for anisotropy, the width of this band (in Å) is computed:

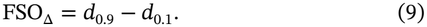

**Notes on** FSO_Δ_

Unlike the other anisotropy metrics in this section, which are dimension-less ratios, FSO_Δ_ is *unbounded* above and has dimensions of length.

## 3. CONICAL FOURIER SHELL CORRELATION AREA RATIO (cFAR)

As in the techniques of 3DFSC and FSO, we compute conical FSCs (cFSCs) to assess directional resolution. For a cone axis 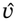 and a fixed cone half-angle *θ*, define the cFSC 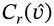 at shell radius *r* as the ordinary FSC (Eq. (2)) restricted to the subset of Fourier voxels on that shell lying within the cone about 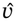:

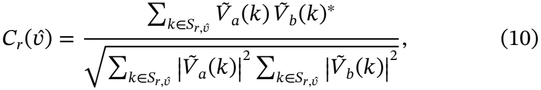

where 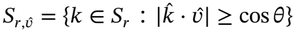 is the set of Fourier voxels on the shell of radius *r* whose direction 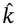 lies within angular distance *θ* of the conical axis ±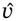.

We compute 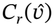 for 3072 directions 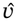 defined by a Fibonacci lattice^22^ on the unit sphere, and for each direction obtain a full curve over *r*. To summarize each curve with a single number, we compute a weighted area under the curve (wAuC), weighting each shell by its surface area 4π*r*^2^ so that higher-frequency shells — which contain proportionally more Fourier voxels — contribute proportionally more to the summary statistic:

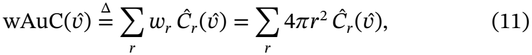

where 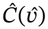 is 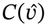 but with all values below 0.143 set to 0.

Fig. 2 shows the output of such a procedure on an example dataset: cFSC curves are coloured by relative wAuC (each direction’s wAuC normalized by the maximum wAuC over all directions). Fig. 2 shows a summary of these cFSCs with statistics computed for each shell (over all conical directions). We define the cFSC Area Ratio (`cFAR`) as the ratio of the worst-to best-performing direction:

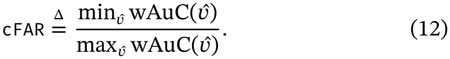

**Figure 2.**
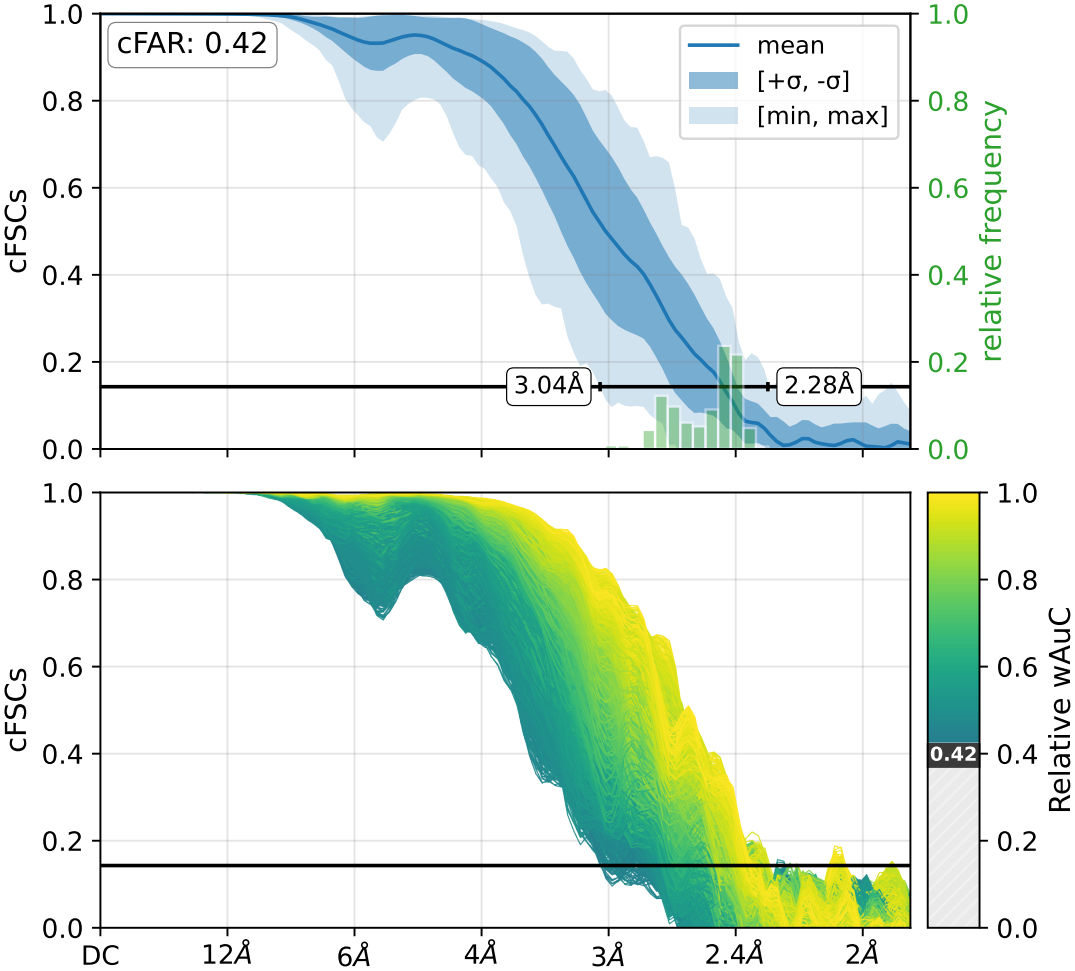
3072 cFSCs from an example dataset (EMPIAR 10668, cf. Table 2) summarized with per-shell statistics, 0.143 crossing histogram and limits and the cFAR score (top). All 3072 cFSC curves coloured by their relative wAuC are shown below, with cFAR identified as the minimum value of relative wAuC (bottom).

Equivalently, `cFAR` is the minimum relative wAuC (visualized on the bottom right in Fig. 2).

**Notes on `cFAR`**

For a perfectly isotropic reconstruction, wAuC is constant and `cFAR` = 1; as resolution becomes more anisotropic, the worst direction’s wAuC falls relative to the best, driving `cFAR` → 0.

## 4. RELATIVE SIGNAL

It is important to note that a conical direction is not the same as a viewing direction: the axis 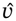 used to define a cone in Eq. (10) does not correspond to the viewing directions that actually contribute signal near 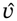. Indeed, recall from Eq. (5) that a particle imaged at viewing direction 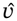 populates the *equatorial* slab of Fourier space orthogonal to 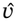. To obtain a diagnostic that correctly reflects the signal contributed by a given viewing direction, we introduce a complementary quantity, the *Relative Signal*, computed over a toroidal region rather than a conical one.

To compute Relative Signal, we consider Fourier-space correlations within a volume akin to a tapered circular slab (Fig. 3). For brevity, we call this a ‘torus,’ noting that, topologically, this volume is not toroidal, as its center is a single point rather than a hole. Concretely, this volume can equivalently be defined as (i) the volume of revolution of a circular sector with central angle 2*ϕ*, or (ii) the spherical *complement* of two antipodal spherical sectors (spherical cones), each with conical half-angle *π*/2 −*ϕ*, as illustrated in Fig. 3.

**Figure 3.**
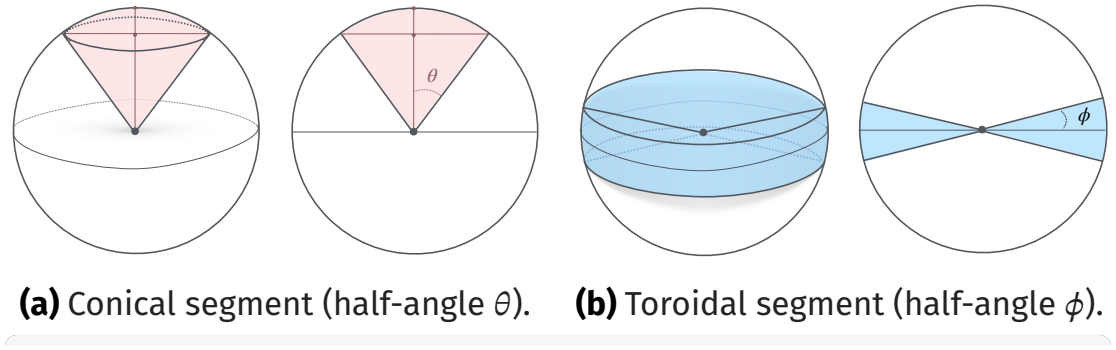
Two different segments of the Fourier sphere used in conical and toroidal FSC calculations.

### 4.1. Cones and Tori

As in Eq. (10), given an axis 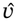, the conical FSC restricts the FSC computation to Fourier voxels 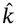 lying within a spherical cone of half-angle *θ* about 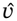:

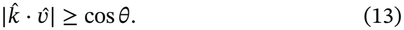

In contrast, given the same axis 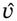, the toroidal FSC restricts the computation to Fourier voxels lying within angle *ϕ* of the equatorial plane orthogonal to 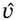 — equivalently, excluding the two antipodal polar caps of half-angle *π*/2 − *ϕ*:

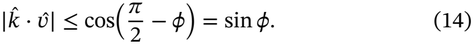

We relate *ϕ* to *θ* by matching the volumes of the two regions (see Appendix A.3 for the full derivation):

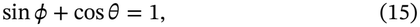

so that, for any conical half-angle *θ* used in the conical FSC, the corresponding toroidal half-width *ϕ* defines a region of matching volume — allowing `cFAR` and its toroidal counterpart to be compared on equal footing. For the default setting of *θ* = 20° (which we use following 3DFSC^8^), *ϕ* ≈ 3.5°.

Analogously to `cFAR`, we define `tFAR` (the toroidal FSC Area Ratio) by replacing the conical FSC *C*_*r*_(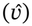) in Eq. (11) with its toroidal counterpart, and taking the same min-over-max ratio across directions 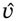:

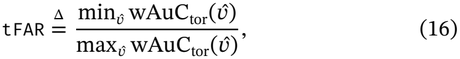

where 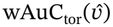 is the weighted area under the toroidal FSC curve (an FSC curve computed within toroidal segments defined by Eq. (14)), and 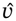 now denotes the toroidal axis (equivalently, the viewing direction for that slab) rather than a conical axis. Because the toroidal region at 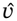 is populated by exactly the particles whose viewing direction lies near 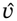, `tFAR` directly measures anisotropy in resolution as a function of *viewing* direction, complementing `cFAR`.

Although `tFAR` can provide an additional anisotropy score, the motivation behind this approach is to find where on the viewing sphere signal is weak by examining 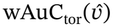. Just as with `cFAR`, we compute the relative wAuC (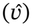, normalized by its maximum over all 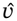) for toroidal segments but now refer to this special quantity as Relative Signal.

Finally, we can now plot Relative Signal as a function of viewing direction. Fig. 4 shows an example of this type of visualization, with Relative Signal shown directly via an atlas, and overlayed in 3D on a rendering of the target structure. These plots let us identify specific views, or clusters of views, that contribute disproportionately weak signal, which the single summary statistics `tFAR` or `cFAR` cannot reveal on their own. To the best of our knowledge, such a method that provides a direct relationship between directional resolution and viewing direction is a novel contribution to the literature.

**Figure 4.**
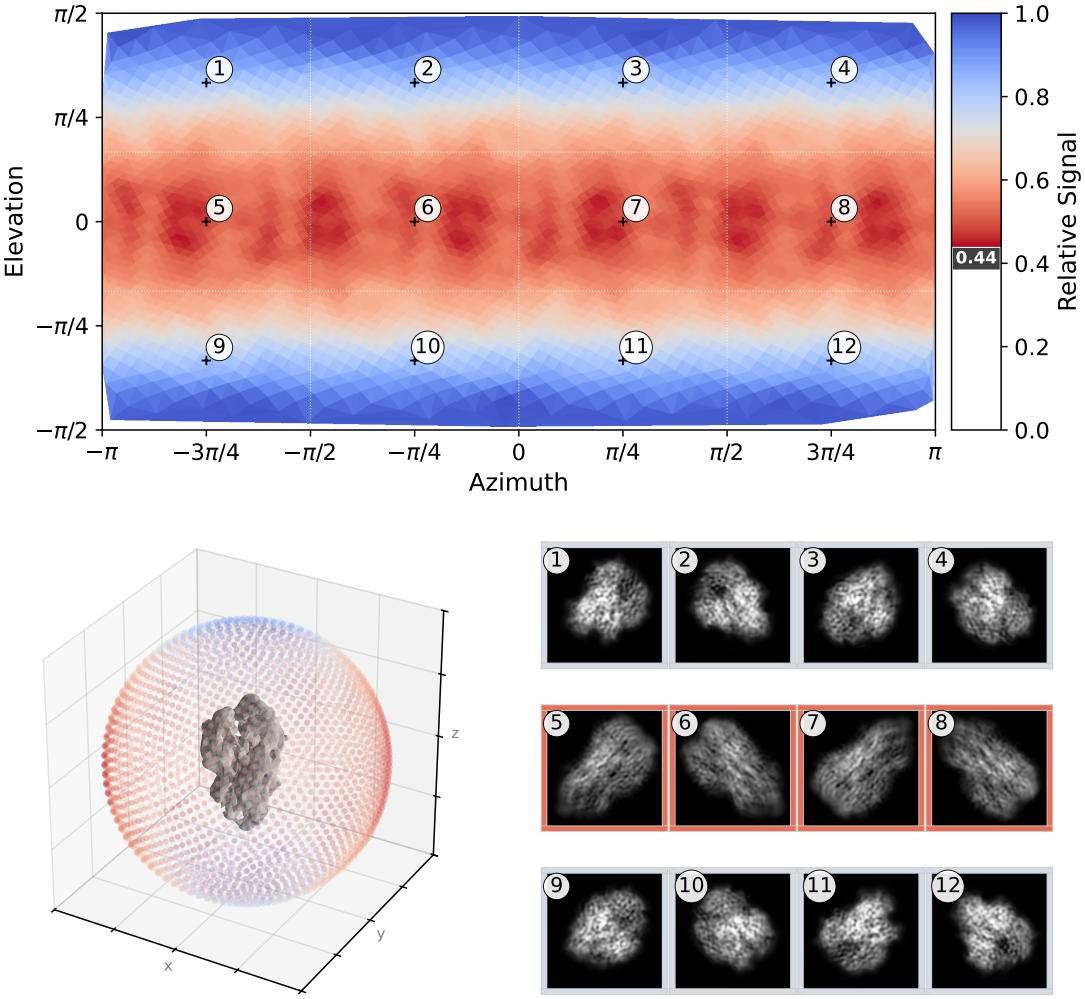
Relative Signal (data from untilted HA trimer, EMPIAR 10096, cf Table 2) visualized against viewing direction (top) with tFAR marked as the minimum value. Each of twelve azimuth/elevation regions is marked at its centroid location. Below, Relative Signal is visualized over the density itself (bottom left), and 12 real-space projections of the density are shown from the viewing directions corresponding to the 12 centroids. Each projection has an outer border whose colour corresponds to the mean Relative Signal in that region, as defined in the upper panel.

## 5. RESULTS AND VALIDATION

We evaluate our approach and compare it to the existing techniques in two complementary settings. First, we apply six orientation diagnostic metrics (including `cFAR` and `tFAR`) to a panel of publicly available cryo-EM datasets chosen to span a wide range of particle sizes, symmetries and orientation distributions. Second, we construct synthetic data in which the orientation distribution of a representative dataset is degraded in two controlled ways, each of which should produce an expected change in the isotropy of the 3D density map. We measure how each orientation metric changes under these degradations to show which metrics successfully capture the degradation. Table 1 summarizes all of the preferred orientation metrics we compare. We refer to the first four (`cFAR`, `tFAR`, FSO_Δ_, Sphericity) as “half-map metrics” because they make use of 3D density maps, and the final two (SCF* and *E*_od_) as “pose-only” to stress that they are functions of particle orientations only. Please refer to the appendix (Table 3) for implementation details of each metric.

**Table 1.**
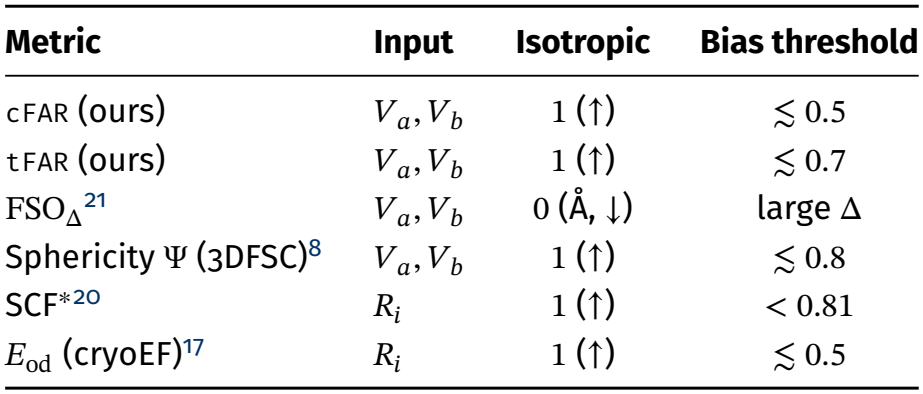
Summary comparison of orientation/anisotropy metrics.

| Metric | Input | Isotropic | Bias threshold |
|---|---|---|---|
| cFAR (ours) | *V<sub>a</sub>*, *V<sub>b</sub>* | 1 (↑) | ≲ 0.5 |
| tFAR (ours) | *V<sub>a</sub>*, *V<sub>b</sub>* | 1 (↑) | ≲ 0.7 |
| FSO<sub>Δ</sub><sup>21</sup> | *V<sub>a</sub>*, *V<sub>b</sub>* | 0 (Å, ↓) | large Δ |
| Sphericity Ψ (3DFSC)<sup>8</sup> | *V<sub>a</sub>*, *V<sub>b</sub>* | 1 (↑) | ≲ 0.8 |
| SCF*<sup>20</sup> | *R<sub>i</sub>* | 1 (↑) | < 0.81 |
| *E*<sub>od</sub> (cryoEF)<sup>17</sup> | *R<sub>i</sub>* | 1 (↑) | ≲ 0.5 |

### 5.1. Experimental cryo-EM data

Our panel of experimental test cases comprises reconstructions from fourteen datasets drawn from the EMPIAR archive, summarized in Table 2. Molecular weights range from 40 kDa to 4000 kDa, particle diameters from 70 Å to 450 Å, box sizes from 192 to 768 pixels, particle counts from *∼* 10^4^ to *∼* 10^6^, and imposed symmetries from C_1_ to O. The panel includes several targets with documented preferred orientation as well as several that are close to isotropic, so that the metrics are exercised across their full operating range.

**Table 2.**
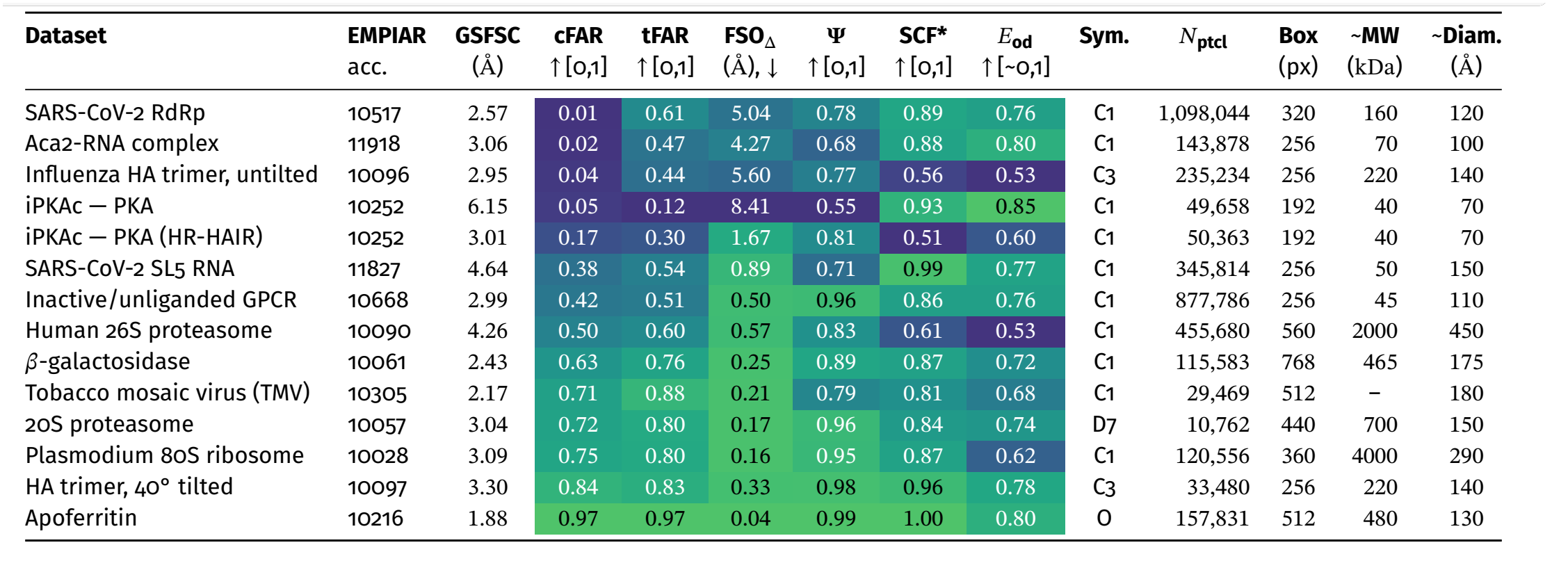
Orientation and anisotropy metrics for the fourteen reconstructions of the real-data panel. Entries are ordered by cFAR, and arrows in the column headings give the direction of improvement. Metric cells are shaded per column on a viridis scale from blue (worst) to green (best) among the datasets shown. Molecular weights and diameters are approximate, and molecular weight is not reported for the TMV filament. EMPIAR-10252 appears twice as two refinements of the same raw data, and the influenza HA trimer appears as an untilted (EMPIAR-10096) and 40° tilted (EMPIAR-10097) acquisition of the same specimen. HR-HAIR refers to the method of *high resolution heterogeneous ab-initio reconstruction*^23^.

| **Dataset** | **EMPIAR** | **GSFSC** | **cFAR** | **tFAR** | **FSO<sub>Δ</sub>** | **Ψ** | **SCF\*** | *E*<sub>od</sub> | **Sym.** | *N*<sub>ptcl</sub> | **Box** | **~MW** | **~Diam.** |
|---|---|---|---|---|---|---|---|---|---|---|---|---|---|
|  | acc. | (Å) | ↑[0,1] | ↑[0,1] | (Å), ↓ | ↑[0,1] | ↑[0,1] | ↑[~0,1] |  |  | (px) | (kDa) | (Å) |
| SARS-CoV-2 RdRp | 10517 | 2.57 | 0.01 | 0.61 | 5.04 | 0.78 | 0.89 | 0.76 | C1 | 1,098,044 | 320 | 160 | 120 |
| Aca2-RNA complex | 11918 | 3.06 | 0.02 | 0.47 | 4.27 | 0.68 | 0.88 | 0.80 | C1 | 143,878 | 256 | 70 | 100 |
| Influenza HA trimer, untilted | 10096 | 2.95 | 0.04 | 0.44 | 5.60 | 0.77 | 0.56 | 0.53 | C3 | 235,234 | 256 | 220 | 140 |
| iPKAc — PKA | 10252 | 6.15 | 0.05 | 0.12 | 8.41 | 0.55 | 0.93 | 0.85 | C1 | 49,658 | 192 | 40 | 70 |
| iPKAc — PKA (HR-HAIR) | 10252 | 3.01 | 0.17 | 0.30 | 1.67 | 0.81 | 0.51 | 0.60 | C1 | 50,363 | 192 | 40 | 70 |
| SARS-CoV-2 SL5 RNA | 11827 | 4.64 | 0.38 | 0.54 | 0.89 | 0.71 | 0.99 | 0.77 | C1 | 345,814 | 256 | 50 | 150 |
| Inactive/unliganded GPCR | 10668 | 2.99 | 0.42 | 0.51 | 0.50 | 0.96 | 0.86 | 0.76 | C1 | 877,786 | 256 | 45 | 110 |
| Human 26S proteasome | 10090 | 4.26 | 0.50 | 0.60 | 0.57 | 0.83 | 0.61 | 0.53 | C1 | 455,680 | 560 | 2000 | 450 |
| *β*-galactosidase | 10061 | 2.43 | 0.63 | 0.76 | 0.25 | 0.89 | 0.87 | 0.72 | C1 | 115,583 | 768 | 465 | 175 |
| Tobacco mosaic virus (TMV) | 10305 | 2.17 | 0.71 | 0.88 | 0.21 | 0.79 | 0.81 | 0.68 | C1 | 29,469 | 512 | – | 180 |
| 20S proteasome | 10057 | 3.04 | 0.72 | 0.80 | 0.17 | 0.96 | 0.84 | 0.74 | D7 | 10,762 | 440 | 700 | 150 |
| Plasmodium 80S ribosome | 10028 | 3.09 | 0.75 | 0.80 | 0.16 | 0.95 | 0.87 | 0.62 | C1 | 120,556 | 360 | 4000 | 290 |
| HA trimer, 40° tilted | 10097 | 3.30 | 0.84 | 0.83 | 0.33 | 0.98 | 0.96 | 0.78 | C3 | 33,480 | 256 | 220 | 140 |
| Apoferritin | 10216 | 1.88 | 0.97 | 0.97 | 0.04 | 0.99 | 1.00 | 0.80 | O | 157,831 | 512 | 480 | 130 |

For every dataset, half-maps and pose estimates were obtained from CryoSPARC^1^ refinement jobs (either homogeneous, non-uniform, or local refinement depending on the workflow used) in v5.0.6. `cFAR` and `tFAR` were computed with a conical half-angle of *θ* = 20° and the volume-matched toroidal half-angle *ϕ* ≈ 3.5° derived above. All metrics are reported in Table 2, where entries are ordered by `cFAR`, and are shown per dataset in Fig. 5. The three lowest (i.e., worst) `cFAR` reconstructions are displayed in Fig. 6 to illustrate the density artifacts present in heavily anisotropic maps.

**Figure 5.**
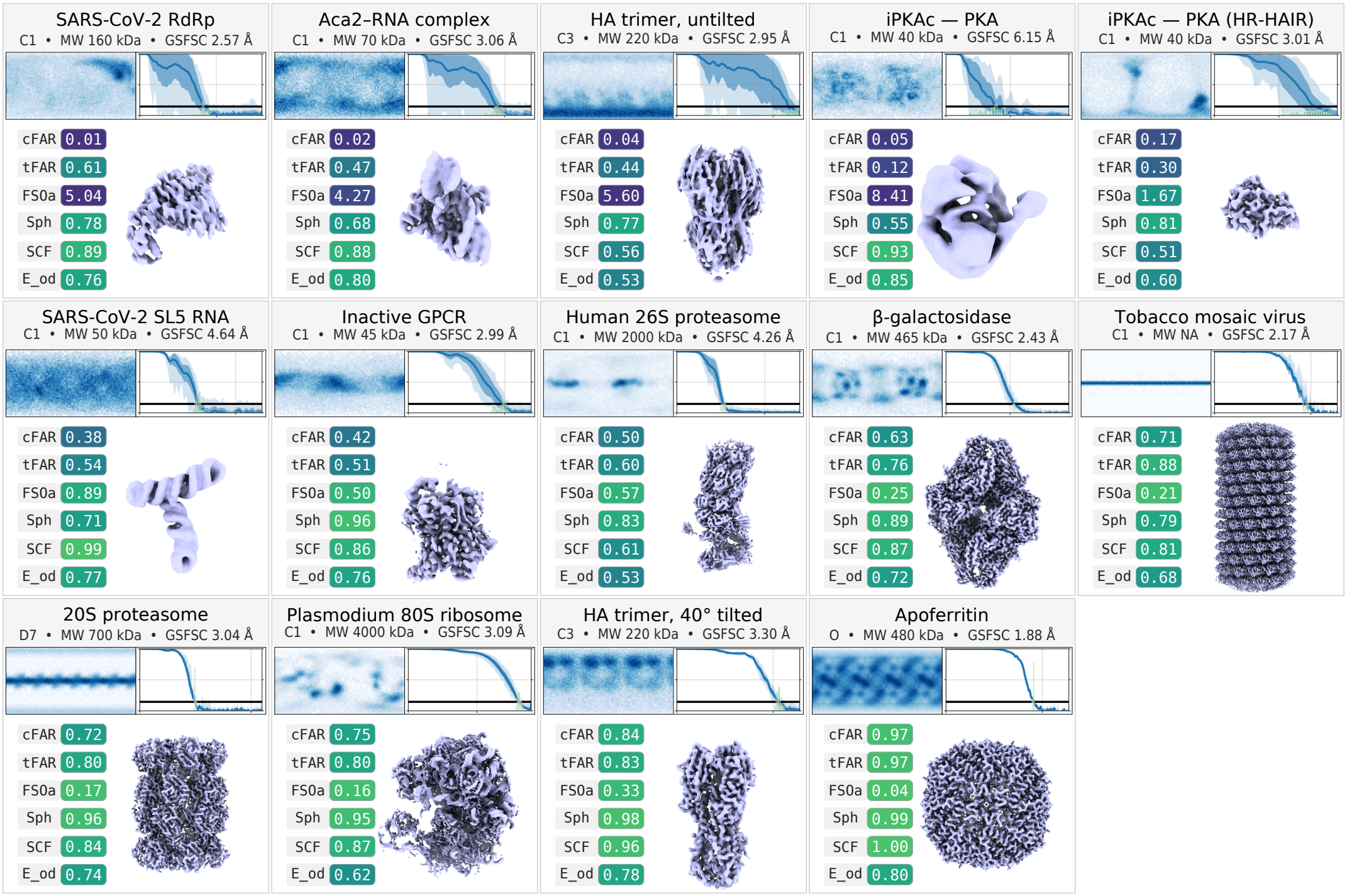
Orientation diagnostics across the real-data panel, ordered by cFAR. Each panel indicates the imposed symmetry, approximate molecular weight and GSFSC resolution. Each panel also displays the computed viewing direction distribution (upper left) and a summary of the cFSC curves (upper right), in which the shaded band reports per-shell statistics over cone directions and the horizontal line marks the 0.143 threshold. Metric values appear beneath, shaded on the same scale as Table 2, alongside a rendering of the corresponding unsharpened density map. The most anisotropic reconstructions have cFSC curves with large variance over cone directions and show visible smearing in the map.

**Figure 6.**
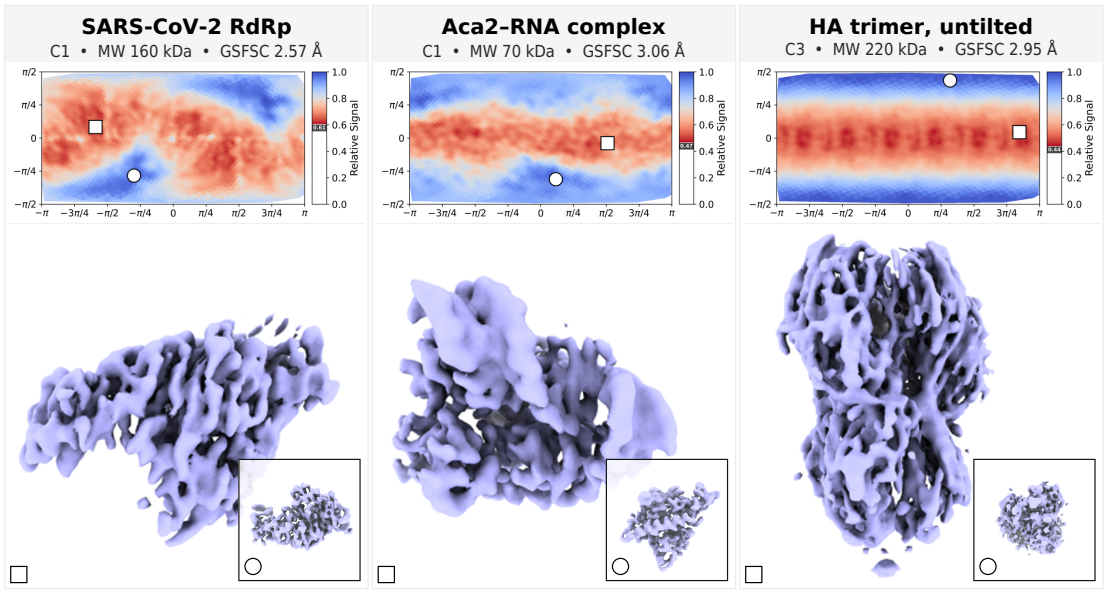
Best and worst views (according to Relative Signal) of the three lowest-cFAR reconstructions in the panel: SARS-CoV-2 RdRp (`cFAR` = 0.01), the Aca2-RNA complex (`cFAR` = 0.02) and the untilted influenza HA trimer (`cFAR` = 0.04). Each map is kept unsharpened and rendered at two different viewing directions: the inset views the map from the direction with the best Relative Signal and the main tile views the map from the direction with the worst Relative Signal. A panel showing Relative Signal as a function of viewing direction is inset above the renders, with the best and worst viewing directions marked with a circle and square, respectively. Note how all three densities report GSFSC resolutions better than *∼*3.1 Å, but the maps do not support this in every direction.

Two entries are deliberately paired with a companion. The EMPIAR-10252 data collection of the iPKAc–PKA sample appears twice, once processed with a standard workflow and once with a gold-standard high-resolution *ab initio* reconstruction workflow^23^, giving two refinements of the same raw data that differ substantially in quality. The influenza HA trimer likewise appears as an untilted (EMPIAR-10096) and a 40° tilted (EMPIAR-10097) acquisition of the same specimen.

Across the panel, `cFAR` scores match well with the level of anisotropy that is visually present in the reconstructed maps. In particular, all maps with clear anisotropy artifacts are scored low by `cFAR`, and those with high `cFAR` do not have anisotropy artifacts. Table 2 is sorted by `cFAR`, and the first four rows correspond to datasets with poor and uninterpretable map quality, yielding `cFAR` scores at or below 0.05. In contrast, on the same datasets, other metrics do not consistently detect the anisotropy; specifically, both pose-only metrics (SCF* and *E*_od_) rank low-quality maps as having high scores.

Overall, `cFAR` has a few properties that make it a useful diagnostic metric in practice. Firstly, `cFAR` is a conservative score, in that across the test set, it does not rate a poor-quality map as having a high score. This means that any map with a low `cFAR` can be automatically and immediately flagged for further user attention. Other metrics may not have the same utility, e.g., Sphericity rates the untilted Influenza HA trimer (with severe anisotropy) almost as high as the Tobacco mosaic virus (with good map quality). `tFAR` is also less conservative than `cFAR` due to toroidal regions having higher angular dispersion than conical regions. Secondly, `cFAR` is highly interpretable: `cFAR` values consistently match with anisotropy across datasets, box sizes, and molecular weights, and it is a bounded metric (zero to one) that makes use of the full range of values, with the 14 test cases spanning 0.01 to 0.97. This means that the severity of anisotropy can be readily understood from the single `cFAR` metric. In contrast, FSO_Δ_ captures the trend of map quality well, but is an unbounded metric and its absolute values have a wide range that is difficult to interpret or threshold. Finally, on the paired datasets, `cFAR` correctly captures the improvement in map quality that is produced by an optimized processing methodology (iPKAc) and by tilting the stage with the same sample (Influenza HA trimer), making it a useful metric to determine the efficacy of such upstream changes.

Results on experimental maps also help illustrate and demonstrate how Relative Signal complements `cFAR`. Fig. 6 shows each of the three worst (by `cFAR` score) maps rendered from the directions with the highest Relative Signal (inset) and lowest Relative Signal (main figure). The well-resolved directions show clear interpretable protein structural features, while the poor-quality directions display smeared or streaking features. Relative Signal therefore helps visualize which particle orientations would be most valuable to collect or curate, in order to improve the isotropy of the 3D maps.

### 5.2. Synthetic Experiments

We construct two series of synthetic experiments in which a single property of an otherwise clean dataset is degraded in controlled increments. Both series begin with a population of particles simulated using a Plasmodium 80S ribosome under the standard image formation model (Eq. (1)). The generating poses are drawn to match the empirical orientation distribution of this dataset, so that the base case is representative of a real orientation distribution (the distribution is visualized in the bottom row, second column from the left of Fig. 5). Quality is then reduced by two orthogonal methods: one alters the content of the particle images while leaving the poses of the real particles intact, and the other alters the poses while leaving the particle images unchanged. In both cases, we expect the isotropy and quality of the resulting 3D reconstructions to become worse, and we measure whether the anisotropy metrics correctly capture this change.

1. **Junk addition**. First, we augment synthetic projections with simulated “junk images” consisting of Gaussian noise together with a small number of randomly placed low-frequency blobs, approximating contaminants encountered in practice. Junk images are assigned poses that are uniform over *SO*(3) and added in increasing amounts to a base set of 50,000 “clean” particles. Fig. 8a shows an example of a clean synthetic image and of a junk image.
2. **Pose jitter**. Second, we perturb, or “jitter”, the poses of a subset of 100,000 clean particles to simulate anisotropic pose accuracy. We select all particles whose pose falls within one quadrant of the viewing direction atlas and perturb their poses by Gaussian noise (i.e., *R*_*i*_ ← *R*_*i*_ exp (*ξ*^×^) with *ξ∼ N*(0, *σ*^2^*I*_3_)) with progressively larger standard deviation (*σ* = 2°, 4°, 6°, 8° and 10°). Fig. 8b illustrates these perturbations using histograms of viewing directions (derived from *R*_*i*_) in blue (clean) and purple (perturbed) colours.

The results of the synthetic experiments are summarized in Fig. 8, which shows means and standard deviations of the different metrics over ten repeats at each increment for both experiments. We omit results for Ψ, the 3DFSC sphericity, for space.

#### 5.2.1. Effects of uniform junk

As illustrated in Fig. 8a, when the viewing sphere is polluted with junk and map quality is degraded, SCF^*^ *improves* monotonically, and the efficiency *E*_od_ remains nearly constant, whereas `cFAR` and `tFAR` (as well as FSO_Δ_) worsen, successfully capturing the presence of uniform junk. Since both `cFAR` and `tFAR` are half-map metrics, they can capture degradations in both particle orientation distributions and the particle quality itself (as is the case here). SCF^*^ and *E*_od_, on the other hand, are primarily functions of *R*_*i*_, so cannot differentiate between clean and junk particles assigned to the same pose.

#### 5.2.2. Effects of anisotropic pose quality

The same pattern is present here (cf. Fig. 8b): SCF^*^ again trends contrary to the change in map quality and *E*_od_ stays flat, whereas `cFAR`, `tFAR`, and FSO_Δ_ worsen as the perturbation variance grows. This series illustrates that our metrics can also capture a systematic degradation to the particle alignments, even if the particle images themselves are clean and the orientations appear to be well-sampled.

### 5.3. Runtime

Finally, we report computational time required to compute each metric in Fig. 7, as recorded during processing of the 14 real datasets in Table 2. For clarity, we sort experiments by box size and symmetry, highlighting the salient dependency for each metric (box size for the half-map metrics and symmetry for the pose-only metrics). Note that, in practice, the runtime of pose-only metrics depends on symmetry (via symmetry expansion), rather than particle count as might be expected, since particles are randomly sub-sampled to a fixed-size subset prior to computation. In all cases, we invested significant effort to ensure each existing metric is computed with reasonable defaults and as efficiently as possible (see Table 3 for details).

**Table 3.**
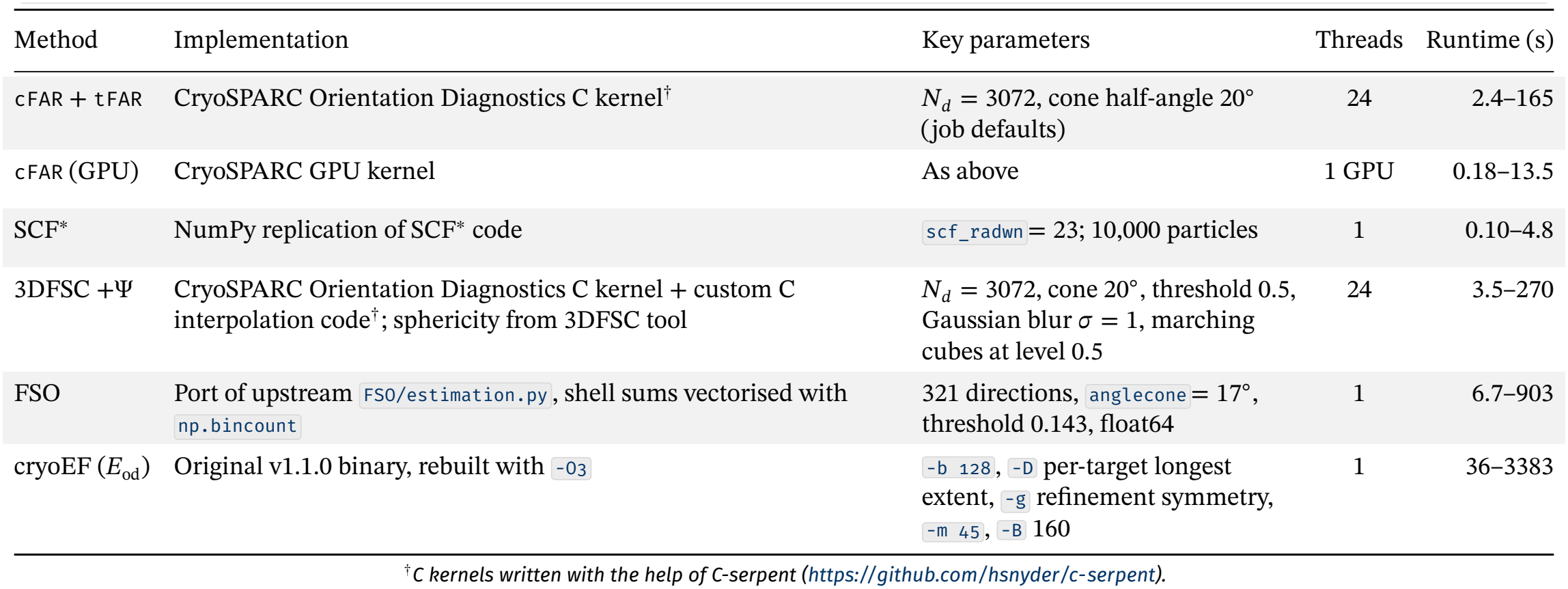
Implementation and runtime details. Thread counts are the values used for the reported timings; runtimes are single runs (*n* = 1) over 14 refinement jobs with box sizes 192–768 px, on a 32-core AMD Ryzen Threadripper PRO 3975WX with an NVIDIA RTX A4000.

| Method | Implementation | Key parameters | Threads | Runtime (s) |
|---|---|---|---|---|
| cFAR + tFAR | CryoSPARC Orientation Diagnostics C kernel<sup>†</sup> | *N<sub>d</sub>* = 3072, cone half-angle 20°<br>(job defaults) | 24 | 2.4–165 |
| cFAR (GPU) | CryoSPARC GPU kernel | As above | 1 GPU | 0.18–13.5 |
| SCF<sup>*</sup> | NumPy replication of SCF<sup>*</sup> code | scf_radwn= 23; 10,000 particles | 1 | 0.10–4.8 |
| 3DFSC +Ψ | CryoSPARC Orientation Diagnostics C kernel + custom C interpolation code<sup>†</sup>; sphericity from 3DFSC tool | *N<sub>d</sub>* = 3072, cone 20°, threshold 0.5, Gaussian blur *σ* = 1, marching cubes at level 0.5 | 24 | 3.5–270 |
| FSO | Port of upstream FSO/estimation.py, shell sums vectorised with np.bincount | 321 directions, anglecone= 17°, threshold 0.143, float64 | 1 | 6.7–903 |
| cryoEF (*E*<sub>od</sub>) | Original v1.1.0 binary, rebuilt with -O3 | -b 128, -D per-target longest extent, -g refinement symmetry, -m 45, -B 160 | 1 | 36–3383 |

<sup>†</sup>*C kernels written with the help of C-serpent (https://github.com/hsnyder/c-serpent).*

**Figure 7.**
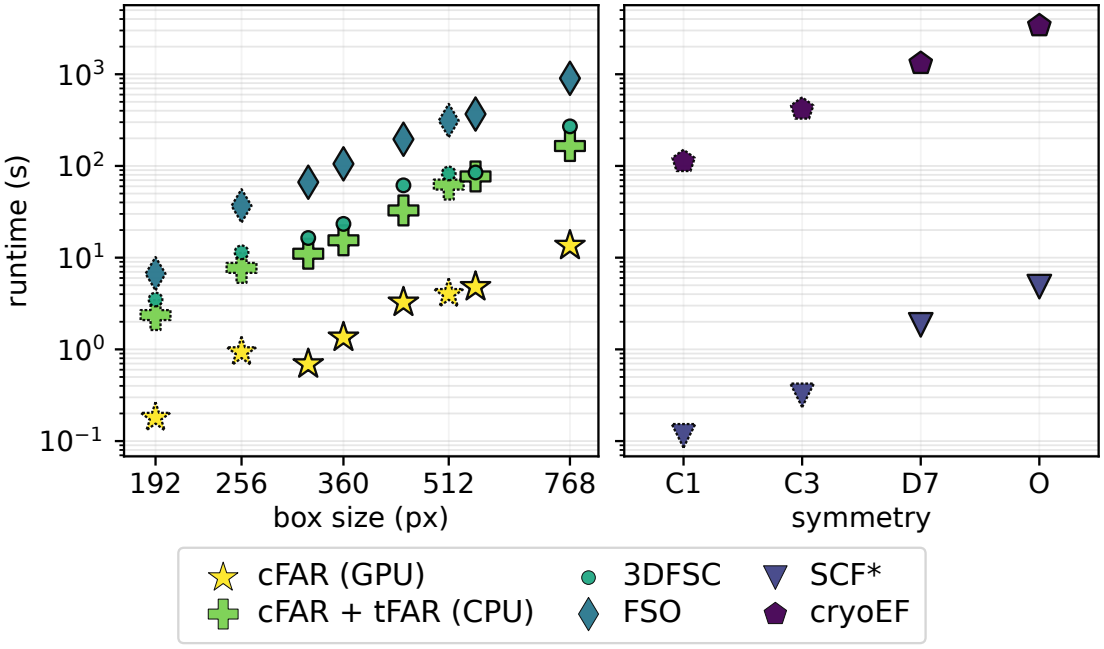
Run-times for the orientation diagnostics on real data, on a semi-log plot. Left: runtime against box size for the half-map-based metrics. Right: runtime against imposed symmetry for the pose-based metrics, whose cost scales with the number of symmetry-expanded poses rather than with grid size (as the number of poses in an asymmetric unit is capped). Markers with dashed edges are arithmetic means over all datasets sharing that box size or symmetry (cf. Table 2).

**Figure 8.**
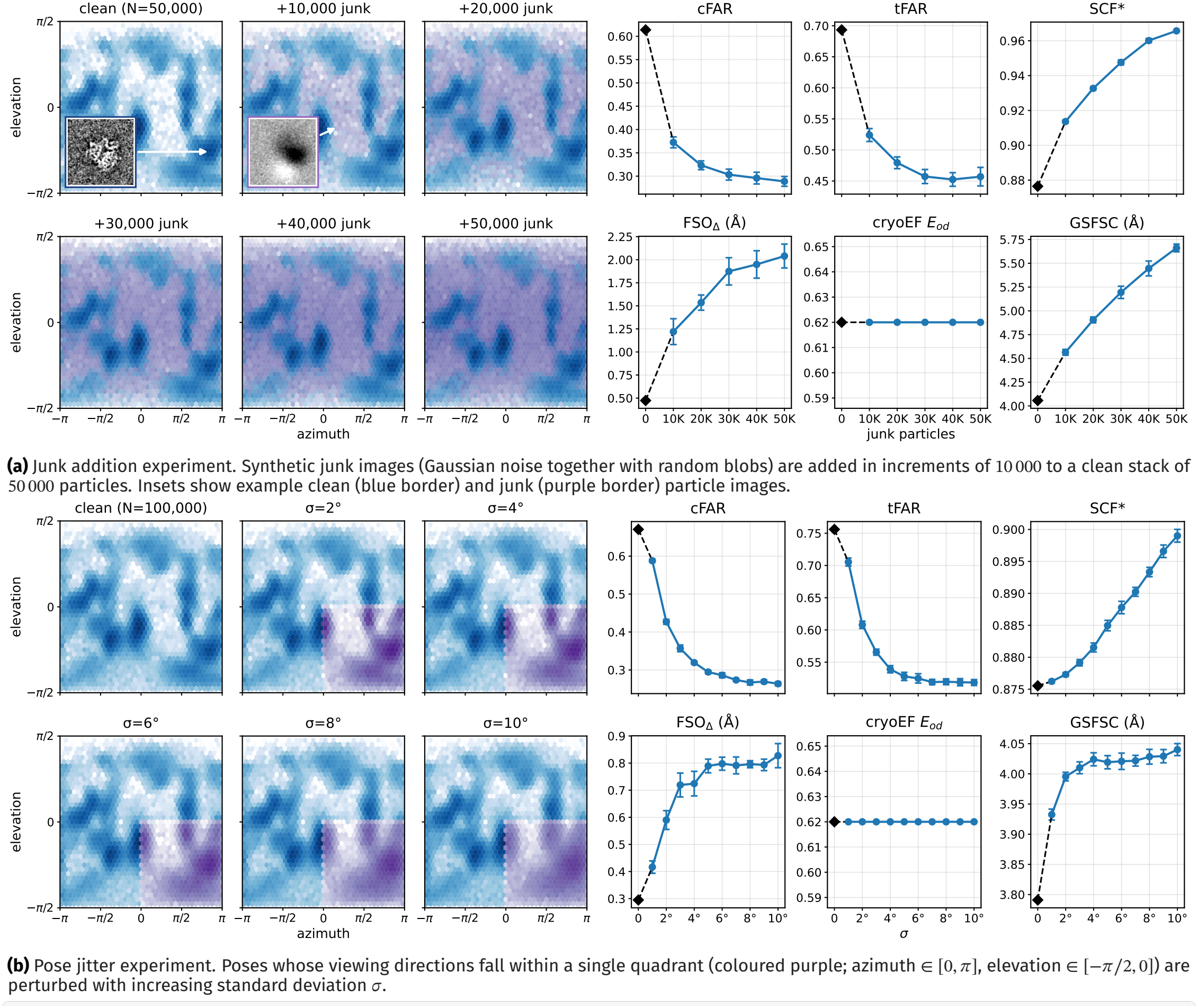
Synthetic degradation series. In each subfigure, the left block shows the assigned viewing direction distribution, and the right block shows the response of five metrics as the degradation increases. Blue bins mark the original particle population and purple bins the degraded component. Black diamonds give the un-degraded base case and are joined to the series by a dashed line; blue markers give the mean over 10 independent replicates with error bars showing standard deviation. Note that the direction of improvement differs between metrics: `cFAR`, `tFAR`, SCF^*^ and *E*_od_ improve upward, while FSO_Δ_ and GSFSC improve downward. In both series GSFSC and all three half-map anisotropy metrics worsen monotonically, whereas SCF^*^ improves and *E*_od_ is unchanged.

Our core `cFAR` and `tFAR` computations use an efficient multi-threaded CPU implementation, and `cFAR` additionally has a GPU implementation built on custom kernels. On most box sizes, the GPU-based `cFAR` computation takes a few seconds or less (using the default set of 3072 conical axes), and the CPU-based `tFAR` and Relative Signal routines finish in under a minute. These runtimes can be an order of magnitude faster than those of the other methods.

## 6. DISCUSSION AND CONCLUSIONS

In this work, we detailed the conical FSC area ratio (`cFAR`), the toroidal FSC area ratio (`tFAR`), and Relative Signal: complementary approaches to deducing the presence of preferred orientation and anisotropy in single particle cryo-EM reconstructions. `cFAR` (and `tFAR`) provide effective numerical summaries of anisotropic resolution, and Relative Signal can identify under-sampled regions of the viewing sphere to help guide upstream efforts to mitigate preferred orientation. We compared and contrasted our methods to four other existing metrics in the literature on real and synthetic data. Across both synthetic degradation series and the fourteen experimental reconstructions, the metrics computed from estimated poses alone did not consistently track map quality, while those computed from half maps, like `cFAR`, did. Furthermore, we illustrated that `cFAR` has a few properties that make it particularly useful as a metric for detecting anisotropy in practice: `cFAR` is a conservative score, and can be used to automatically flag reconstruction results; `cFAR` is highly interpretable, being consistent, bounded (zero to one), and sensitive across the full range of values, so that the severity of anisotropy can be easily understood; and `cFAR` correctly captures the improvement in map quality resulting from upstream changes in processing or data collection. Despite these useful properties, any single number metric must be interpreted with care. `cFAR` and `tFAR` inherit many of the caveats of FSC-based analysis: they require a completed refinement, and they respond to masking and to imposed symmetry, which raises apparent directional signal without adding independent information. Both are also extremum-based statistics, reporting on a single worst direction rather than an average; thus plots that show full directional distributions (e.g., Relative Signal) should be consulted for a more comprehensive analysis. In general, none of the metrics presented in this work can completely replace the essential step of *visually examining* a density map in 3D, nor are they intended to.

The methods presented in this work are included in the CryoSPARC software suite, with efficient CPU and GPU implementations, for free academic use.

## 7. COMPETING INTERESTS

CryoSPARC™ is developed and distributed by Structura Biotechnology Inc.

## 8. DATA AND SOFTWARE AVAILABILITY

All data processing in this work was carried out using CryoSPARC v5.0.6, available at cryosparc.com. All raw data was downloaded from EMPIAR^5^; see accession codes in Table 2.

## 9. ACKNOWLEDGMENTS

We thank the entire team at Structura Biotechnology Inc. that designs, develops, distributes, maintains and supports the CryoSPARC software system with which this project was implemented and tested. In particular, we thank Saara Virani for her help in editing and polishing early drafts of this work. We also thank all contributors to the EMPIAR data repository.

## 10. ADDITIONAL DETAILS

### 10.2. Typesetting

This document was typeset in LATEX, with an adapted class derived from rho class.

## A. APPENDICES

### A.1 Implementation Details

Please refer to Table 3 for parameters and implementation details for all metrics.

### A.2 Full timings

Please refer to Fig. 9 for runtimes for each individual real dataset.

**Figure 9.**
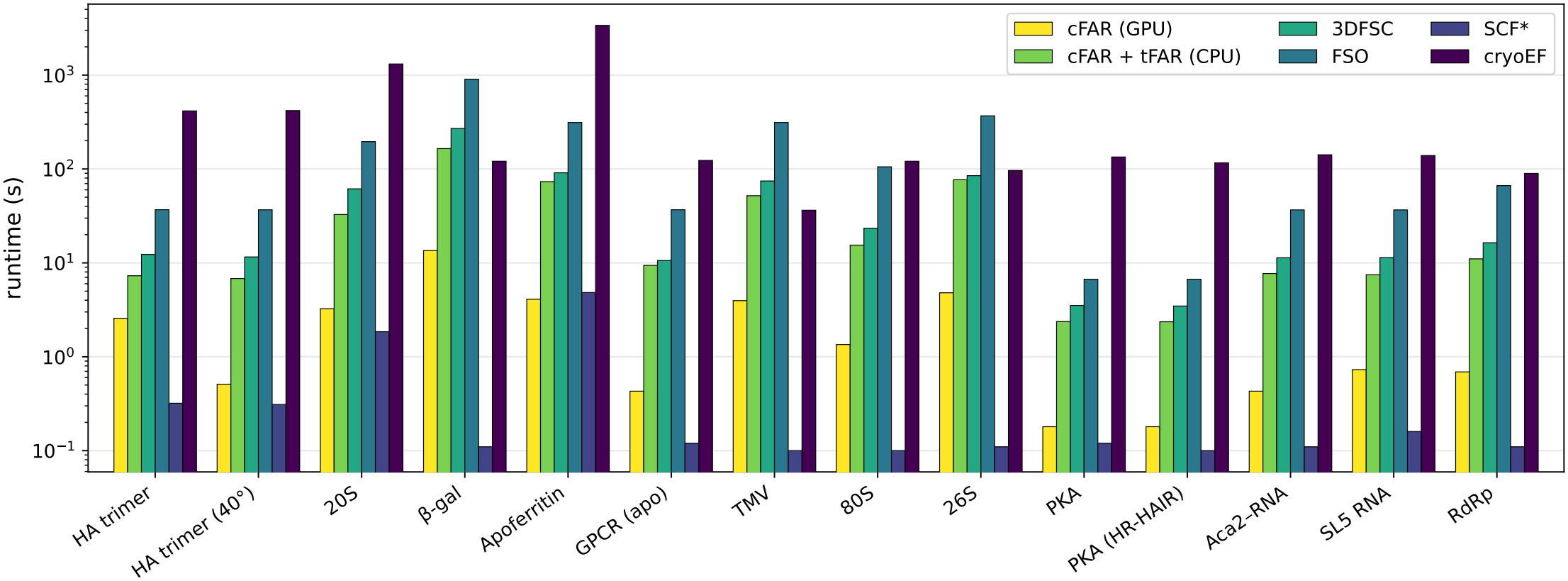
Runtimes for six implementations of five metrics across the fourteen different datasets tested. The GPU-based implementation of `cFAR` is an order of magnitude quicker than other half-map metrics.

### A.3 Setting *ϕ* by Matching Volumes

With each conical FSC, we consider a Fourier volume *V*_*c*_ comprised of two spherical cones (arranged anti-podally to account for Hermitian symmetry):

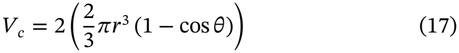

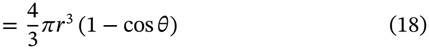

The volume of the ‘torus’, *V*_*t*_, can be derived via the complement-based definition:

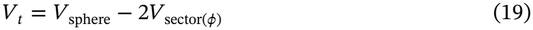

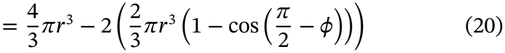

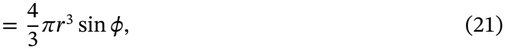

where we have used the fact that 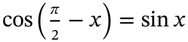. Setting *V* = *V*_*t*_, we arrive at the simple relation:

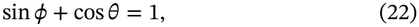

from which we can derive the toroidal half-angle *ϕ*, given a conical half-angle *θ*.

## Notes

https://www.cryosparc.com

### Summary of Updates

Revised for layout and formatting; no content changes.

